# Loss of Rnf31 and Vps4b sensitizes pancreatic cancer to T cell-mediated killing

**DOI:** 10.1101/2021.07.29.453937

**Authors:** Nina Frey, Luigi Tortola, David Egli, Sharan Janjuha, Kim Fabiano Marquart, Tanja Rothgangl, Franziska Ampenberger, Manfred Kopf, Gerald Schwank

## Abstract

Pancreatic ductal adenocarcinoma (PDA) is an inherently immune cell deprived tumor, characterized by desmoplastic stroma and suppressive immune cells. Here we systematically dissected PDA intrinsic mechanisms of immune evasion by *in vitro* and *in vivo* CRISPR screening, and identified Rnf31 and Vps4b as essential factors required for escaping CD8^+^ T cell-killing. Using murine PDA cells and human PDA organoids, we demonstrate that *Rnf31* protects from TNF-mediated caspase 8 cleavage and subsequent apoptosis induction. For *Vps4b* we found that inactivation impairs autophagy, resulting in increased accumulation of CD8^+^ T cell-derived granzyme B and subsequent tumor cell lysis. Orthotopic transplantation of Rnf31− or Vps4b deficient pancreatic tumors, moreover, revealed increased CD8^+^ T cell infiltration and effector function, and markedly reduced tumor growth in mice. Our work uncovers vulnerabilities in PDA that might be exploited to render these tumors more susceptible to the immune system.

## Introduction

Immune evasion is a common trait of most human cancers. Through phenotypic changes tumor cells evade recognition of effector T cells and modulate the tumor microenvironment to establish an immune suppressive niche ^1,2^. While immune checkpoint inhibition shows great potential for curative cancer treatment, pancreatic ductal adenocarcinoma (PDA) is largely refractory to immunotherapy ^3,4^. Among the described mechanisms responsible for the highly effective immune evasion of PDA are (i) insufficient antigenicity ^5^, (ii) high expression of PD-L1 ^6^, (iii) exclusion of dendritic cells while attracting T regulatory cells ^2^ and suppressive myeloid populations ^7,8^, and (iv) the sequestration of major histocompatibility complex class 1 (MHC-I) ^9^. In order to better understand cell-autonomous mechanisms that protect tumors from immune clearance, genome-wide CRISPR-Cas9 screens have been performed in melanoma, renal-, colorectal- and breast cancer cell lines. Together they identified the interferon-ɣ (IFNɣ) response, TNF-mediated NF*κ*B signaling and autophagy as core pathways involved in immune evasion across different cancer types ^10–18^. However, to our knowledge, a comprehensive genetic analysis of potential target genes to enhance anti-tumor immunity in PDA is still missing.

Here we used genome-wide *in vitro* CRISPR screening and targeted *in vivo* CRISPR screening to systematically reveal positive and negative regulators of cytotoxic T lymphocyte (CTL) sensitivity in PDA. In addition to previously described genes involved in the regulation CTL-mediated tumor cell killing, we identify *Rnf31* and *Vps4b* as central components for PDA immune escape *in vitro* and *in vivo*. Our results suggest that *Rnf31,* as part of the linear ubiquitination chain assembly complex (LUBAC), mediates immune-escape by stabilizing anti-apoptotic proteins in the TNF pathway, and that *Vps4b,* as part of the autophagy machinery, reduces susceptibility to T-cell mediated tumor cell lysis by lowering intracellular granzyme B contents. The elucidated mechanisms of immune evasion in PDA provide potential strategies for enhancing efficacy of cancer immunotherapies.

## Results

### A genome-wide CRISPR screen identifies regulators of immune evasion in PDA

To identify genes modulating CTL-mediated killing of PDA we performed a pooled, genome-wide CRISPR knock-out screen in pancreatic cancer cells. We first engineered a PDA cell line derived from the autochthonous KPC mouse model (***K****ras*^G12D^, *Tr****p****53*^R172H/+,^ *Pdx*-**C**re), which stably expresses *Sp*Cas9 and chicken ovalbumin (OVA). Cells were subsequently transduced with a murine single-guide (sg)RNA library targeting 19 647 genes at a 500x coverage ^19^. To mimic cytotoxic T cell killing, we co-cultured cancer cells for three days with activated, OVA-specific CD8^+^ T cells (OT-I T cells), followed by a three-day recovery period prior to DNA isolation for analysis by next generation sequencing (NGS) from the surviving cell population [Fig. 1a]. Validating screening conditions and sufficient library representation, we observed a strong overlap of depleted sgRNAs targeting essential genes in OT-I T cell treated- and untreated KPC cells [Supplementary Figs. 1a, b]. Next, we inspected differentially distributed sgRNAs, and defined genes targeted by enriched sgRNAs as resistors and genes targeted by depleted sgRNAs as sensitizers for CTL-mediated killing (FDR < 0.1). We identified several genes with a well-characterized role in CTL-mediated killing in different cancer types, demonstrating that the previously described core cancer intrinsic CTL evasion gene network is also conserved in PDA [Figs. 1b, c] ^10–12,14,20^. For example, genes associated with the IFNɣ pathway (*Jak1*, *Jak2*, *Ifngr1*, *Ifngr2*, *Stat1)* and antigen presentation machinery (*B2m*, *Tap1*) conferred resistance to CTL-mediated PDA killing upon inactivation [Figs. 1c, d, Supplementary Fig. 1c], and genes regulating TNF-triggered apoptosis (*Cflar, Traf2*), NF*κ*B signaling (*Nfkbia*, *Tnfaip3*) and autophagy (*Atg5*, *Atg7, Atg10, Atg12*, *Gabarapl2*) sensitized PDA cells to CTL-mediated killing upon inactivation [Figs. 1c, d, Supplementary Fig. 1c]. Interestingly, the two strongest sensitizers to T cell-mediated killing identified in our PDA screen were *Rnf31* and *Vps4b,* but for both genes mechanistic insights in context of CTL sensitivity are lacking.

**Figure 1:**
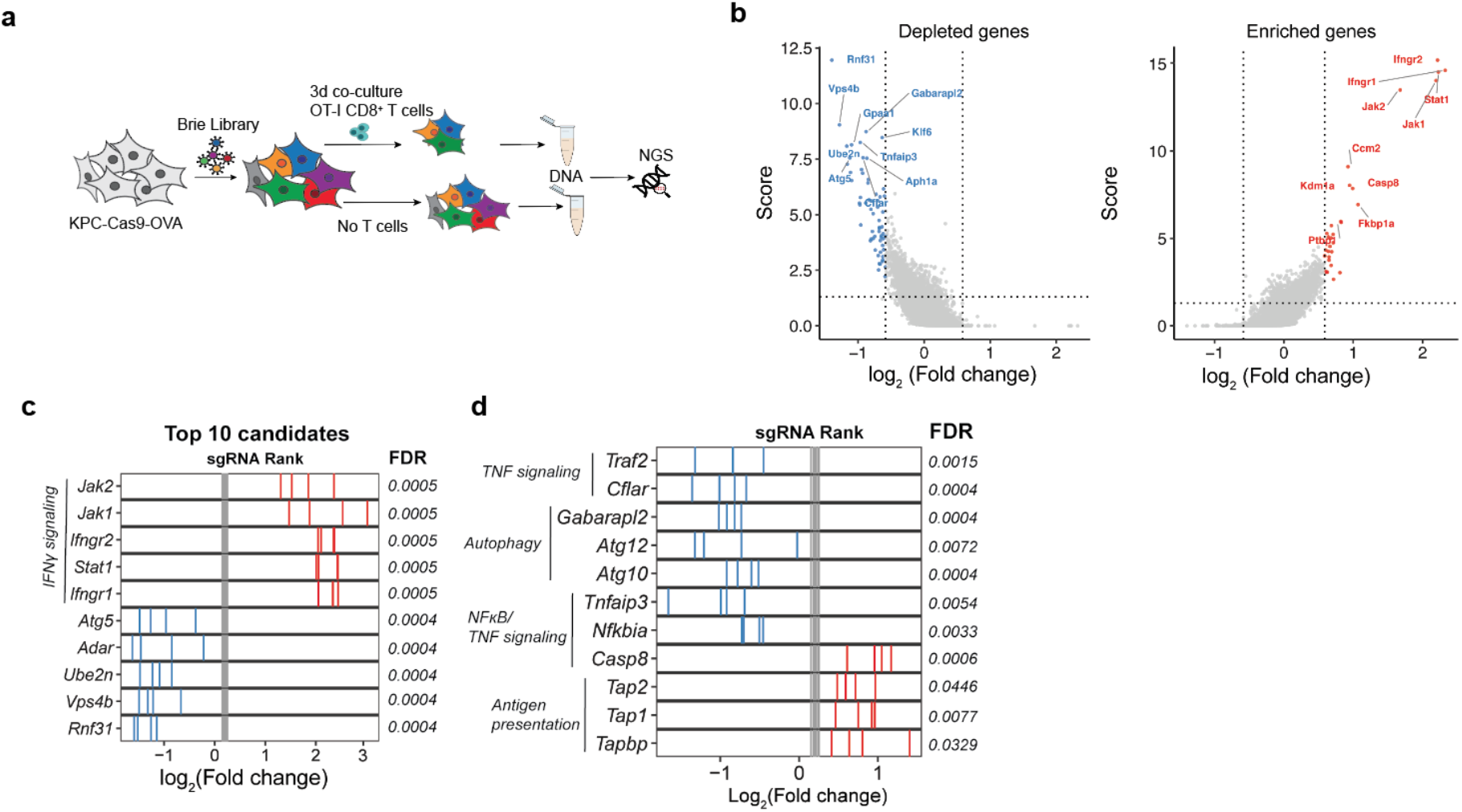
Genome-wide CRISPR screen in PDA cells reveals immune evasion mechanisms in vitro. **(a)** Schematic of genome-wide in vitro CRISPR screen. **(b)** Volcano plot of top ten depleted (blue) and enriched (red) genes. Screening analysis was performed with MaGeCK RRA. **(c)** sgRANK of the top (red) and bottom (blue) five depleted genes is represented. Grey bars represent non-targeting sgRNAs. **(d)** sgRANK of the enriched (red) and depleted (blue) genes of different immune evasion pathways. Grey bars represent non-targeting sgRNAs.

### A targeted CRISPR screen validates immune modulators *in vivo*

To explore whether top candidates from the *in vitro* screen also affect CTL-mediated PDA killing *in vivo* we next performed a targeted library screen in mice. We generated a secondary library targeting 63 genes (hits with FDR < 0.1) with ten sgRNAs per gene, and containing 600 non-targeting control sgRNAs as well as seven positive control sgRNAs targeting ovalbumin. We furthermore omitted several IFNɣ pathway components to avoid redundancy. The library was transduced into KPC-Cas9-OVA cells, which we subsequently orthotopically transplanted into pancreata of RAG1^−/−^ mice. After tumor formation, we adoptively transferred activated CD8^+^ OT-I T cells to tumor-bearing mice and collected the residual tumors five days later [Fig. 2a]. In vivo validated candidates showed consistent phenotypes across all mice [Fig. 2b], and in line with previous studies we observed a substantial, albeit not complete overlap between *in vitro* and *in vivo* screening results [Fig. 2c, Supplementary Fig. 2a] ^21^. Resistors of CTL evasion included well-known immune evasion genes, such as *Stat1* and *Casp8,* as well as the positive control *ovalbumin* [Fig. 2b]. Among sensitizers of CTL killing - which are of particular therapeutic interest as they bear the potential to enhance anti-tumor immunity in PDA upon inhibition - were the previously described genes *Adar* and *Cflar* ^22–24^, as well as the two strongest sensitizers identified in our CRISPR screen, *Vps4b* and *Rnf31* [Figs. 2b, d], prompting us to further study their role in PDA immune evasion.

**Figure 2:**
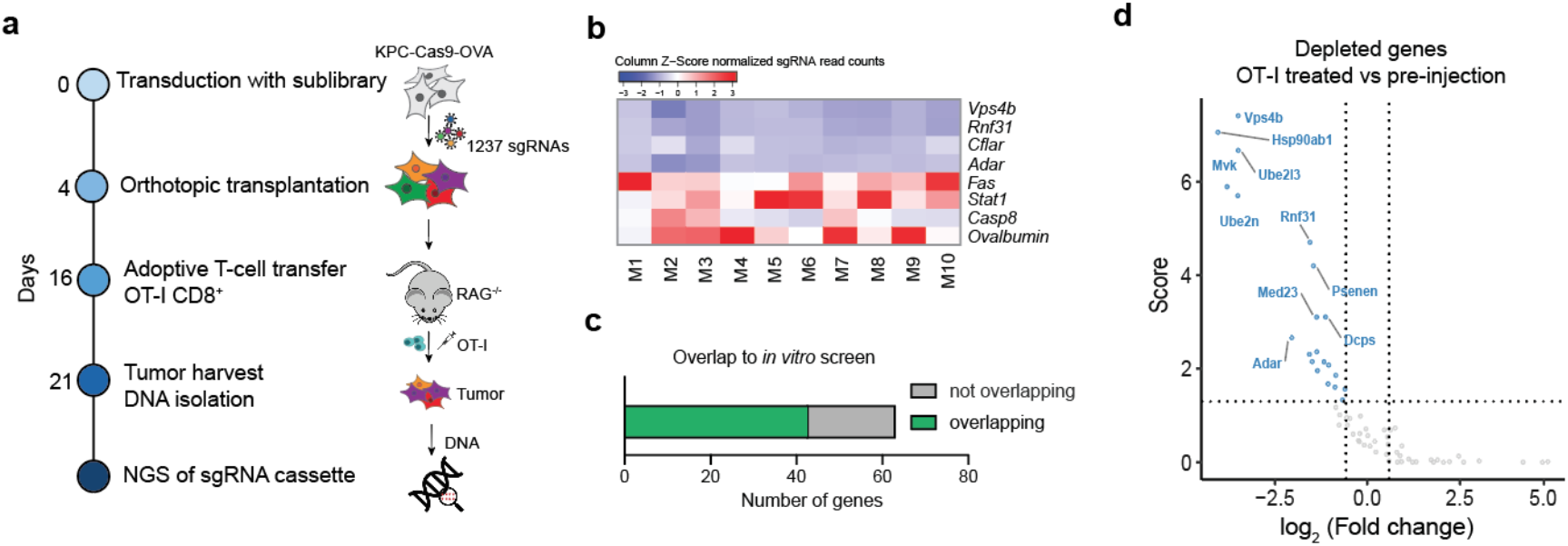
A targeted CRISPR library screen validates candidates in vivo. **(a)** Schematic of the secondary CRISPR screen *in vivo*. (**b**) Heatmap of normalized read counts of sgRNAs across ten individual mice (M1 - M10). **(c)** Bar diagram of sublibrary genes in comparison to their predicted phenotype (from *in vitro* screen). **(d)** Volcano plot of depleted (blue dots) genes of the sublibrary *in vivo* screen when comparing the pool of preinjected KPC cells to OT-I CD8^+^ T cell treated tumors.

### A competition assay confirms the role of *Rnf31* and *Vps4b* in immune evasion

We then performed arrayed validation of *Vps4b-* and *Rnf31*-mediated PDA sensitization to CTL killing in a competition assay. KPC-Cas9-OVA cells carrying *Vps4b*- and *Rnf31-* targeting sgRNAs were labeled with mCherry^+^, and KPC-Cas9-OVA cells carrying a non-targeting control sgRNA were labeled with GFP^+^. After confirming that knock-outs do not affect cell proliferation per-se [Supplementary Fig. 3a], we co-cultured candidate lines with CD8^+^ OT-I T cells and assessed mCherry and GFP proportions by flow cytometry [Fig. 3a]. As expected, without generating a gene knock-out in mCherry^+^ cells we did not observe a shift in the mCherry:GFP ratio after the addition of OT-I T cells [Figs. 3b, c], while targeting *Stat1,* a well-known resistor of CTL killing and positive control for our assay, shifted the ratio towards IFNɣ- signaling deficient mCherry^+^ cells [Figs. 3b, c]. In contrast, *Vps4b*^KO^ and *Rnf31*^KO^ KPC cells had a strong growth disadvantage under immune attack, leading to an increase of the GFP^+^ control cell population [Figs. 3b, c]. Our results therefore confirm that inhibition of *Rnf31* and *Vps4b* sensitizes PDA to CTL killing.

**Figure 3:**
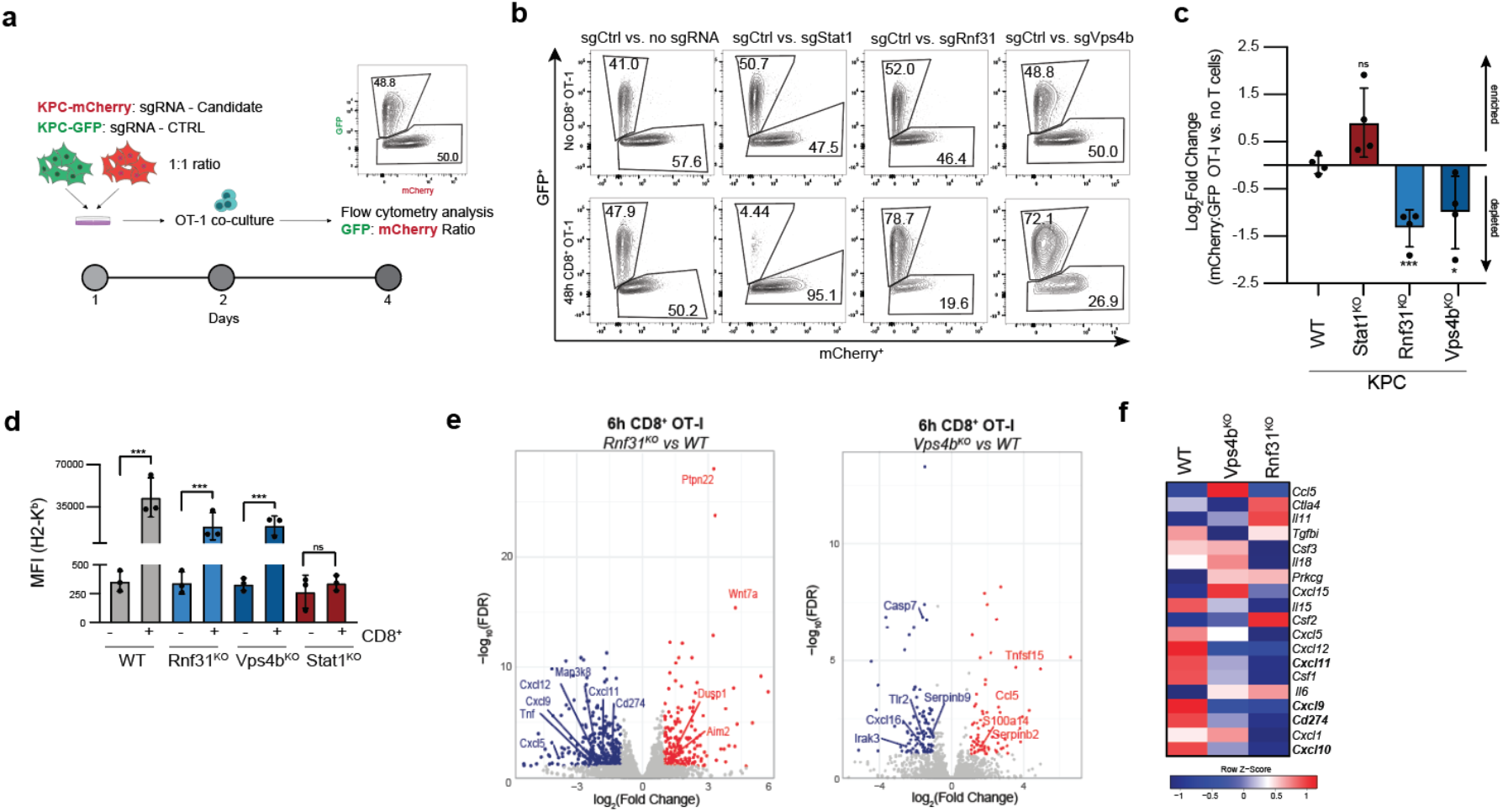
Arrayed validation of selected screening hits *in vitro*. **(a)** Schematic of *in vitro* competition assay. **(b)** Representative flow cytometry plots (GFP vs. mCherry) of the arrayed hit validation with and without T cell co-culture. **(c)** Quantification of **(b)**. Log2 Fold change of the mCherry:GFP ratio OT-I treated compared to the matched untreated condition. **(d)** Mean fluorescence intensity (MFI) of Pan-H2-K^b^. (**e)** Volcano plots of differentially expressed (DE) genes in *Rnf31*^KO^ and *Vps4b*^KO^ cells after 6h of OT-I T cell exposure compared to equivalently treated KPC^WT^ cells. Highlighted genes are putatively involved in anti-tumor immunity. DE genes in red/blue: | Log2FC | > 1, FDR < 0.1. **(f)** Heatmap of normalized counts per million (CPM) of selected immune modulatory factors after OT-I T cell exposure across different genotypes. Significance in **(c)** was determined with an unpaired two-tailed t test. Significance in **(d)** was determined with one-way ANOVA analysis. *p < 0.05, **p < 0.01, ***p < 0.001; ns, non-significant, p > 0.05. Values represent mean ± SD, data are derived from at least three independent experiments.

### Functional characterization of the role of *Rnf31* and *Vps4b* in immune evasion

Antigen presentation by major histocompatibility complex I (MHC-I) proteins is necessary for efficient anti-tumor immunity. As shown in a recent study, pancreatic cancer cells commonly sequestrate MHC-I to evade the adaptive immune system ^9^, prompting us to suspected that loss of *Rnf31* and *Vps4b* facilitates CD8^+^ mediated killing by increasing MHC-I levels on PDA. To test this hypothesis, we assessed surface MHC-I levels in KPC cells upon exposure to CTLs. Confirming our assay, we observed robust induction of MHC-I upregulation in parental KPC cells, which, as expected, was perturbed in IFNɣ signaling deficient *Stat1*^KO^ cells (Fig. 3d, Supplementary Fig. 3d). Next, we analyzed MHC-I induction in *Rnf31*^KO^ and *Vps4b*^KO^ KPC cells. However, we observed similar surface MHC-I levels compared to the parental cell line (Fig. 3d), demonstrating that enhanced CTL-mediated killing is not triggered by an increase in antigen presentation [Fig. 3d].

Next, we sought to gain insights into the transcriptional networks mediating the sensitizing effects of *Rnf31*^KO^ and *Vps4b*^KO^ to CTL killing. We therefore performed RNA-sequencing (RNA-seq) on the different PDA knock-out lines with and without six hours of CD8^+^ T cell exposure [Supplementary Fig. 3b]. Confirming functional gene knock-outs, transcript levels of *Rnf31* and *Vps4b* were downregulated in the respective PDA lines [Supplementary Fig. 3c]. Furthermore, *Rnf31*^KO^ and *Vps4b*^KO^ cell lines displayed relatively mild, but consistent transcriptional changes [Fig. 3e]. Among the differentially expressed genes in CTL-treated *Rnf31* or *Vps4b* knock-out PDA cells were several cytokines and chemokines, including the Cxcr3 ligands *Cxcl9/10/11* [Figs. 3e, f]. Notably, the Cxcr3-Stat3 signaling axis has previously been described to enhance PDA aggressiveness and contribute to an immune suppressive environment through inducing PD-L1 (CD274) expression ^25,26^, and low expression of these chemokines is correlated with a better prognosis in human PDA patients [Supplementary Fig. 3e]. We therefore hypothesize that downregulation of Cxcr3 ligands contributes to the immune stimulatory environment triggered by *Rnf31* and *Vps4b* inactivation.

### Rnf31 loss sensitizes PDA to TNF- induced apoptosis via caspase 8

Cytotoxic T cells induce death in target cells via different processes, including the release of TNF, secretion of granules filled with granzymes and perforins, and by engaging the Fas-FasL axis. To systematically explore which of these effector mechanisms are sensitized upon *Rnf31* and *Vps4b* inhibition, we first assessed tumor cell sensitivity to TNF ligands. Interestingly, we found that parental KPC cells and *Vps4b*^KO^ KPC cells were insensitive to TNF-induced apoptosis, but that *Rnf31*^KO^ KPC cells rapidly underwent cell death upon TNF treatment [Fig. 4a]. Engagement of the TNF receptor triggers several signaling branches, including pro-survival NF*κ*B signaling as well as apoptosis induction via caspase 8 cleavage ^27–29^. Notably, *Rnf31* has previously been reported to function as an E3 ubiquitin-protein ligase within the linear ubiquitination chain assembly complex (LUBAC), which is involved in regulating NF*κ*B signaling and in stabilizing anti-apoptotic proteins such as c-Flip ^27^. We therefore speculated that the *Rnf31* knock-out sensitizes tumor cells to TNF-mediated apoptosis either indirectly, by abrogating NF*κ*B pro-survival signaling, or directly, by facilitating caspase 8 cleavage. When we first assessed TNF-mediated NF*κ*B activation, we found phosphorylation of the NF*κ*B subunit p65/Rela in all genetic backgrounds, including *Rnf31*^KO^ cells [Fig. 4b], indicating functional NF*κ*B signaling. When we next analyzed caspase 8 cleavage upon TNF treatment, activation was observed in *Rnf31*^KO^ KPC cells but not in parental KPC- or *Vps4b*^KO^ KPC cells [Fig. 4b]. Hence, our data suggest that in Rnf31-deficient cells intact NF*κ*B signaling is not sufficient to rescue TNF-activated caspase 8 cleavage.

**Figure 4:**
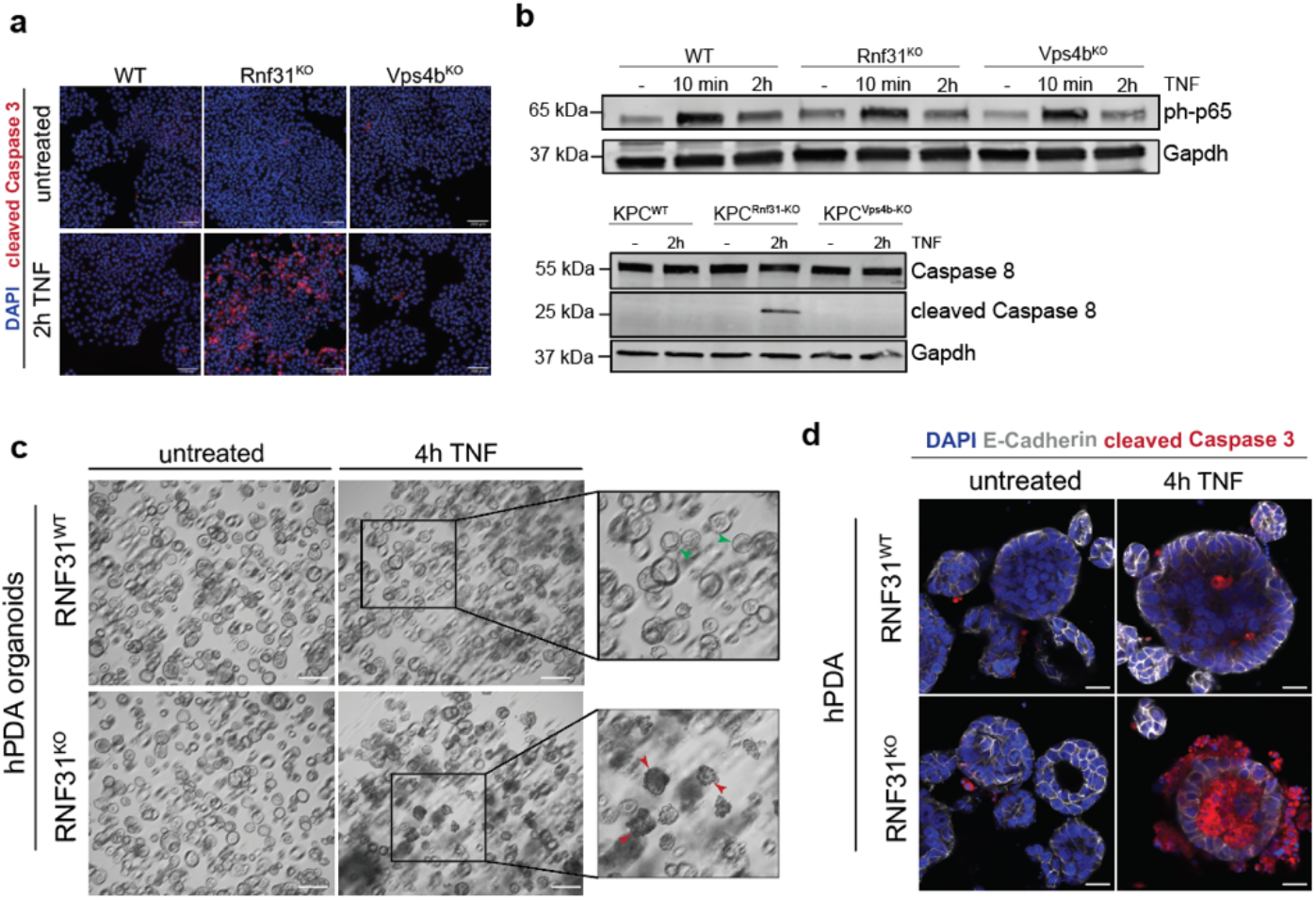
*Rnf31*^KO^ sensitizes PDA to TNF-triggered apoptosis. **(a)** Immunofluorescence staining of KPC candidate lines after 2h of 100 ng/ml TNF. Cleaved caspase 3 (red) and DAPI (blue). Scale bar represents 100 *μ*m. **(b)** Western Blot analysis of KPC cell lines after TNF treatment (100 ng/ml) for active NF*κ*B signaling (phospho-p65; upper panel) and cleaved caspase 8/caspase 8 (bottom panel). Gapdh was included as loading control. **(c)** Brightfield images of human PDA organoids in the presence of 100 ng/ml TNF for 4h. Boxes highlight viable (green arrows) and dying organoids (red arrows). Scale bar represents 200 *μ*m. **(d)** Whole mount staining of human PDA organoids after 4h TNF (100 ng/ml) treatment with cleaved caspase 3 (red), E-cadherin (white) and DAPI (blue). Scale bar represents 20 *μ*m.

To further assess whether loss of Rnf31 also sensitizes human PDA to TNF-mediated cell death, we next generated patient-derived and engineered human pancreatic cancer organoids (hPDA) with *RNF31*^KO^ mutations, and treated these organoids for four hours with TNF. In line with results from murine PDA tissues, only *RNF31*^KO^ but not *RNF31*^WT^ PDA organoids activated apoptotic cell death upon TNF stimulation [Figs. 4c, d, Supplementary Figs. 4a, b]. Taken together, our results suggest that loss of the LUBAC subunit Rnf31 sensitizes murine and human pancreatic cancer to CTL killing by rendering cells susceptible to caspase-8-mediated apoptosis upon TNF signaling [Supplementary Fig. 4c].

### Vps4b depletion impairs functional autophagy and increases intracellular Granzyme B levels

As part of endosomal sorting complexes required for transport III (ESCRT-III) Vps4b functions as an AAA-type ATPase involved in diverse processes regulating protein homeostasis, including the catalyzation of phagophore closure during autophagy ^30^. Considering that several autophagy-related genes have been identified as sensitizers for CTL-mediated killing, we reasoned that *Vps4b*^KO^ cells might sensitize PDA to CTL-killing by inhibiting autophagy. To test this hypothesis, we transduced cells with an autophagic flux reporter, and assessed if autophagy is impaired in *Vps4b* knock-out KPC cells ^31^. The reporter consists of a LC3-GFP-LC3ΔG-RFP fusion protein; LC3-GFP is localized to the autophagosome and degraded during autophagy, and LC3ΔG-RFP lacks a C-terminal glycine and stably resides in the cytoplasm during autophagy [Fig. 5a]. While parental KPC cells showed a strong upregulation of autophagy upon starvation [Fig. 5b], *Vps4b*^KO^ KPC cells showed an impaired autophagic flux, similar to fully autophagy-deficient *Atg5*^KO^ KPC cells [Fig. 5b]. These data suggest that loss of Vps4b sensitizes PDA to CTL killing through disrupting autophagy.

**Figure 5:**
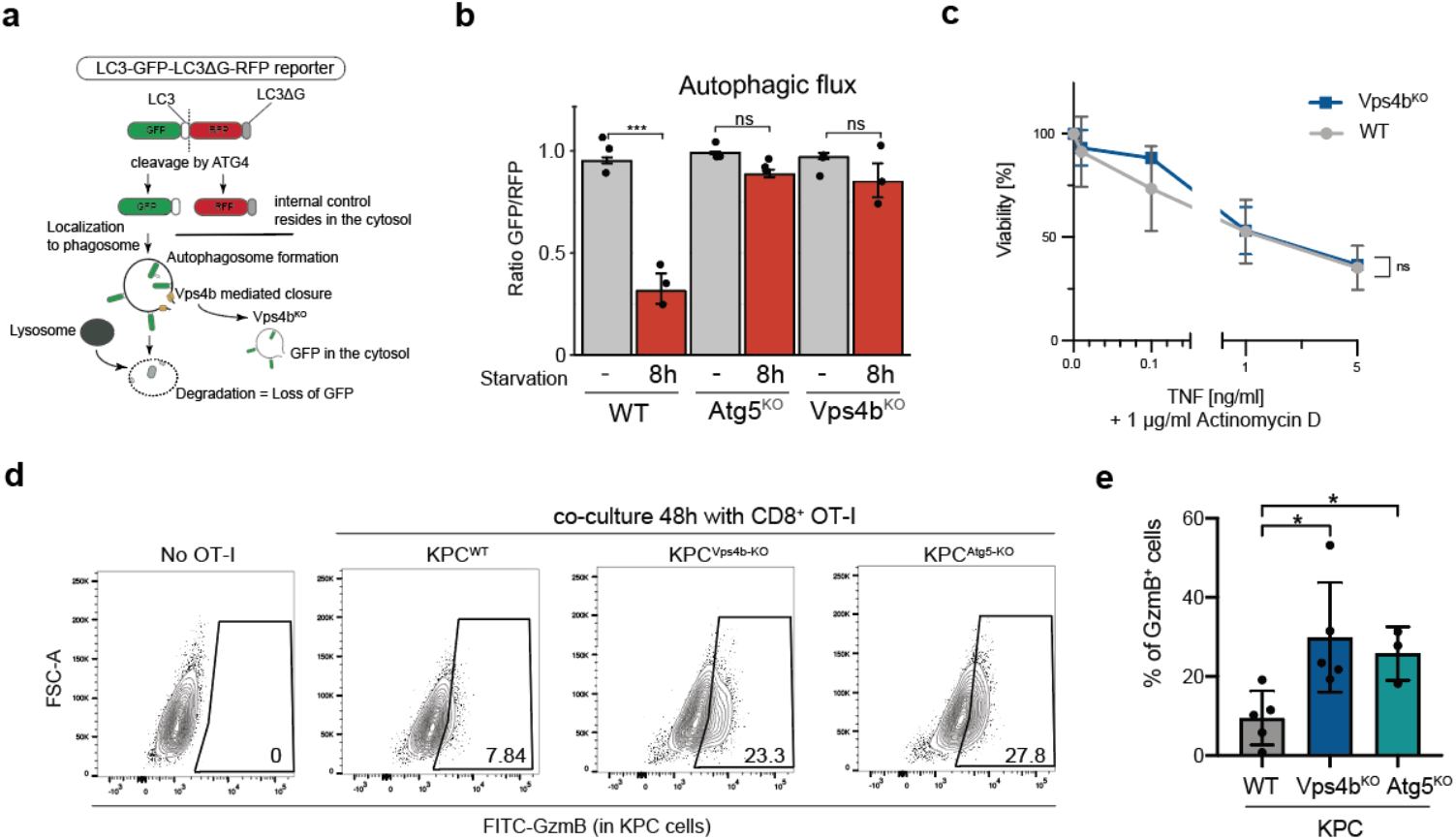
Disruption of Vps4b leads to impaired autophagy and granzyme B accumulation in tumor cells. **(a)** Schematic of autophagic flux reporter based on the LC3-GFP-LC3ΔG-RFP probe adapted from Kaizuka *et al.* ^31^. **(b)** Quantification of autophagic flux by flow cytometry in different KPC lines under normal and starvation conditions (8h in PBS + 2% FBS). Bars represent the ratio of GFP to RFP expressing cells. **(c)** Assessment of TNF sensitivity threshold in WT and *Vps4b*^KO^ KPC cells in the presence of 1 *μ*g/ml actinomycin D. Crystal violet staining was used for quantification of viable cells. Represented data is relative to untreated control cells. (**d**) Flow cytometric analysis of intracellular granzyme B in KPC cells. Cells were gated on FSC/SCC – viability - CD8- negative. **(e)** Quantification of granzyme B positive cancer cells and T cells based on **(d)**. Significance in **(c)** was determined with an unpaired two-tailed t test. Significance in **(b)** was determined with one-way ANOVA. Significance in **(e)** was determined with an unpaired, two-tailed t-test. *p < 0.05, **p < 0.01, ***p < 0.001; ns, non-significant, p > 0.05. Values represent mean ± SD, data are derived from at least three independent experiments.

In a previous study it was suggested that high autophagy rates in PDA contribute to immune evasion by sequestering surface MHC-I levels ^9^. In KPC cells, nevertheless, we observed a robust induction of MHC-I surface expression upon CTL exposure, which was also not affected by Vps4b depletion (Fig. 3d). In another model it was suggested that autophagy inhibition facilitates CTL-mediated killing through increasing sensitivity to TNF-induced cell death ^11^. However, we did not observe apoptosis induction when we treated Vps4b^KO^ KPC cells with TNF [Fig. 4a]. In addition, when we sensitized KPC cells to TNF-induced apoptosis through Actinomycin D - a transcriptional inhibitor of pro-survival NF*κ*B signaling - prior to TNF treatment, *Vps4b*^KO^ KPC cells were again not sensitized to increasing TNF concentrations compared to parental KPC cells [Fig. 5c]. Interestingly, a recent studies found that high autophagy levels in breast cancer cells promote NK cell-derived granzyme B degradation, resulting in resistance to cytotoxic immune cells ^32,33^. We therefore hypothesized that autophagy deficiency could sensitize PDA cells to CTL killing through insufficient granzyme B clearance. Hence, we quantified intracellular granzyme B levels in KPC cells upon OT-I T cell exposure. Indeed, while OT-I T cells produced comparable amounts of granzyme B, autophagy deficient *Atg5*^KO^- and *Vps4b*^KO^ KPC cells accumulated more granzyme B compared to parental KPC cells [Figs. 5d, e, Supplementary Fig. 5a]. Taken together, our data suggest that *Vps4b* inhibition perturbs autophagy and thereby reduces the capability of PDA cells to degrade granzyme B upon CTL-mediated killing. Importantly, this phenotype was not only limited to *Vps4b*-deficient cells, but generally linked to PDA cells with impaired autophagy, providing a model how autophagy influences sensitivity to CTL-mediated killing.

### *Rnf31* and *Vps4b* inhibition increases CTL infiltration and effector function in vivo

To further characterize increased CTL susceptibility of *Rnf31*^KO^ and *Vps4b*^KO^ KPC cells *in vivo*, we next analyzed the effect of these mutations on PDA progression and tumor microenvironment in mice. Therefore, we orthotopically transplanted KPC cells with different genotypes (wildtype, *Rnf31*^KO^ and *Vps4b*^KO^) into C56BL/6 animals, and assessed survival and tumor weight, as well as immune cell composition and effector function using flow cytometry [Fig. 6a, Supplementary Fig. 6d]. While remaining Cas9 expression in these lines did not affect tumor growth [Supplementary Fig. 6a], loss of *Rnf31* and *Vps4b* markedly decreased tumor mass and resulted in significantly enhanced survival of tumor-bearing mice [Fig. 6b]. In case of *Vps4b*^KO^ tumors the effect was strongly dependent on adaptive immunity, since tumor mass reduction was not apparent in RAG1^−/−^ mice [Supplementary Fig. 6b]. *Rnf31*^KO^ tumors, however, also showed reduced growth compared to KPC^WT^ tumors in RAG1-deficient hosts, most likely due to the continued expression of TNF and other death receptor ligands by NK cells [Supplementary Fig. 6b]. We next analyzed immune cell infiltration and CD8^+^ T cell effector function across the different tumor genotypes. While loss of *Vps4b* and *Rnf31* did not cause significant changes in macrophages, CD11c^+^ dendritic cells, CD4^+^ T helper cells, NK cells and CD4^+^ Foxp3^+^ regulatory T cells, we detected a minor increase of neutrophils in *Rnf31*^KO^ tumors [Fig. 6c, Supplementary Fig. 6c]. In addition, we observed a substantial increase of infiltrating CD8^+^ T cells in *Vps4b*^KO^ and *Rnf31*^KO^ tumors compared to parental KPC tumors [Fig. 6c]. Further analysis of CD8^+^ CTL markers for effector function revealed a significant reduction in exhausted PD1^+^ CD8^+^ T cells in *Vps4b*^KO^ and *Rnf31*^KO^ tumors [Fig. 6d], concomitant with an increase in cytokine production; in *Rnf31*^KO^ tumors TNF production and in *Vps4b*^KO^ tumors TNF and IFNɣ production was increased in infiltrating CD8^+^ T cells [Fig. 6d]. Together, these findings demonstrate that loss of *Rnf31* and *Vps4b* sensitize PDA to CTL-mediated killing also in a non-cell-autonomous manner, through increasing CTL effector function, thereby feeding into a forward loop with cell-autonomous mechanisms to enhance anti-tumor immunity.

**Figure 6:**
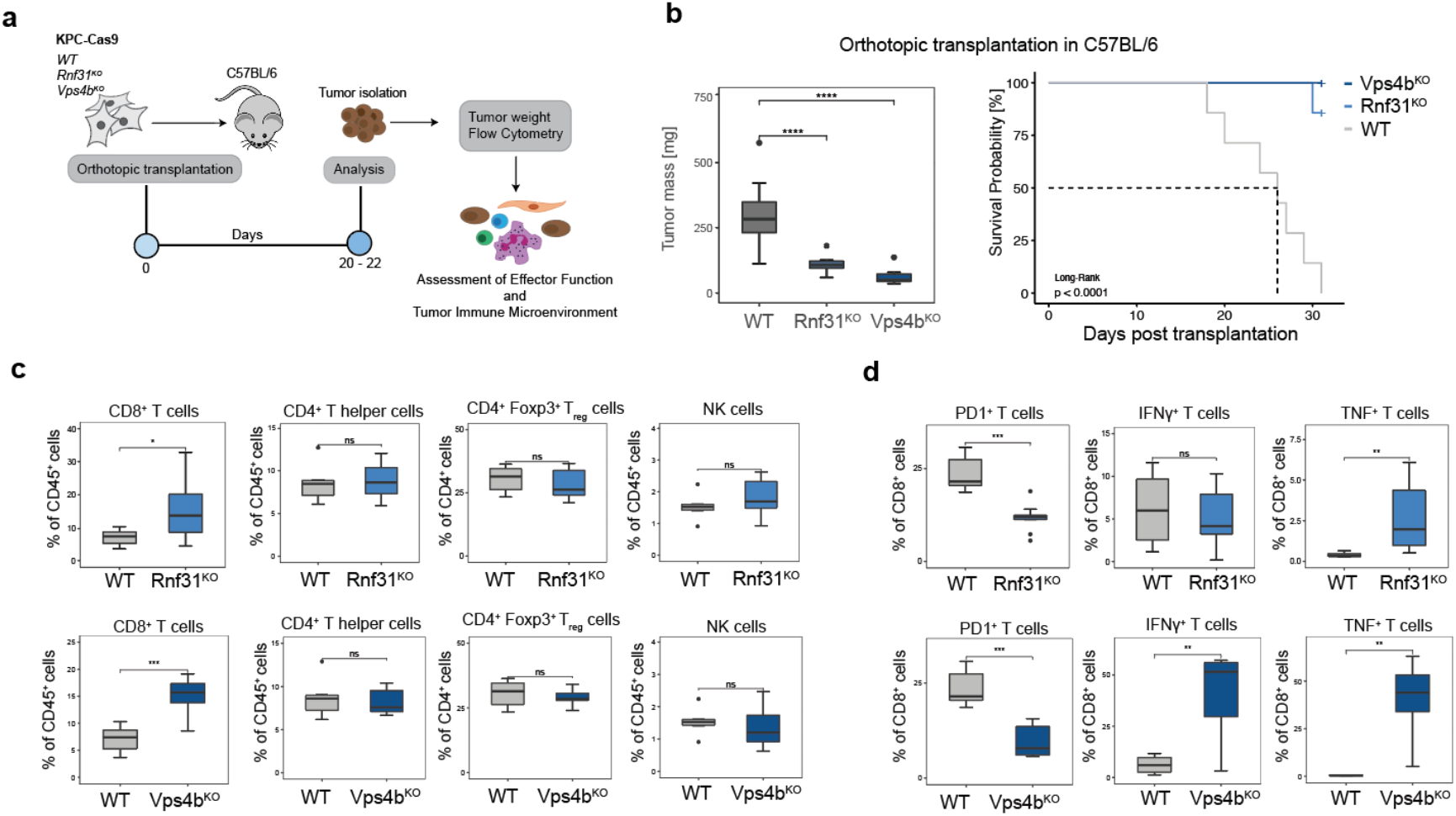
*Rnf31*^KO^ and *Vps4b*^KO^ enhances CD8^+^ T cell function *in vivo*. **(a)** Schematic of in vivo experimental set up. **(b)** Tumor weight and survival after orthotopic transplantation into C5BL/6 mice. **(c)** Flow cytometry analysis of immune cell population within tumors. **(d)** Flow cytometry analysis of effector function of CD8^+^ T cells within tumors. Significance in **(b, panel 1)**, **(c)** and **(d)** was determined with an unpaired two-tailed t test. *p < 0.05, **p < 0.01, ***p < 0.001; ns, non-significant, p > 0.05. The middle line in the boxplots shows the median, the lower and upper hinges represent the first and third quartiles, and whiskers represent ±1.5× the interquartile range.

## Discussion

Several recent studies performed CRISPR screening in PDA to study metastasis formation, metabolic vulnerabilities, combinatorial drug targeting and therapy resistance ^21,34–36^. Here, we applied *in vitro* and *in vivo* CRISPR screening in PDA to interrogate tumor intrinsic mechanisms of immune evasion. One of the strongest sensitizers to CTL-mediated killing was *Rnf31,* for which we show that its inactivation facilitates TNF-induced apoptosis via caspase 8 cleavage. Notably, previous work has already linked TNF resistance to immune evasion ^11,37^. However, these studies have been conducted in a TNF-susceptible colorectal cancer cell line (MC38), impeding the identification of TNF sensitizers such as *Rnf31*. In contrast, KPC pancreatic cancer cells are intrinsically resistant to TNF, which allowed us to unravel a mechanism that abates TNF resistance.

Another strong sensitizer to CTL-mediated killing identified in our screen was *Vps4b,* which could be linked to autophagy. Autophagy has recently been postulated as an important modulator of anti-tumor immunity in several cancer entities ^9–11,38^. However, in contrast to previous findings we did not observe increased MHC-I antigen presentation ^9^ or enhanced TNF-induced apoptosis ^10,11^ upon autophagy inhibition. Instead, we provide evidence that impaired autophagy leads to reduced granzyme B clearance, suggesting that *Vps4b* or *Atg5* depletion facilitates tumor cell lysis by CD8^+^ T cells through enhanced granzyme B accumulation.

Taken together, we used functional genomics approaches to identify mechanisms for circumventing immune evasion in PDA. Analysis of two of the strongest hits, *Vps4b* and *Rnf31,* demonstrated that their inhibition sensitizes tumor cell clearance directly, via cell-autonomous mechanisms, and indirectly, by increasing the number and functionality of intertumoral CD8^+^ T cells. Our insights in sensitizing pancreatic cancer to the host immune system could open up novel strategies to enhance the efficacy of T cell-mediated tumor killing, potentially allowing PDA patients to benefit from the vast advances made in the field of cancer immunotherapy in the future.

## Methods

### Animals

Wildtype C57/BL6 mice were obtained from Charles River Laboratories. RAG1^−/−^ (NOD.129S7(B6)-*Rag1*^tm1Mom^/J) and OT-I (C57BL/6-Tg(*TcraTcrb*)1100Mjb/J) were obtained from Jackson Laboratories and bred in-house. All animals were housed in a pathogen-free animal facility in cages with up to five animals at the Institute of Molecular Health Sciences at ETH Zurich and kept in a temperature- and humidity-controlled room on a 12h light–dark cycle. All animal experiments were performed in accordance with protocols approved by the Kantonales Veterinäramt Zurich in compliance with all relevant ethical regulations.

### Cell culture

The KPC cell line (C57/BL6 background) was generated by Dr. Jen Morton (Beatson Institute) and purchased at Ximbio (Cat# 153474). KPC cells were derived from primary KPC tumors obtained from Pdx-Cre; Kras^G12D/+^; Trp53^R172H/+^ mice. All KPC lines used in this study were cultured in Iscove’s Modified Dulbecco’s Medium (IMDM, 31980030, Gibco) supplemented with 10% fetal bovine serum (FBS), 1% penicillin/streptomycin (Gibco) and 50 *μ*M β-Mercaptoethanol (Gibco). Cells were incubated at 37°C in 5% CO_2_. The parental KPC line was engineered with Lenti-Cas9-Hygromycin and Lenti-Ovalbumin-mCherry-Blasticidin constructs (for details see “Plasmids”) in order to express Cas9 and full-length Ovalbumin (KPC-Cas9-OVA).

### Plasmids

For generation of Lenti-EF1α-Cas9-P2A-HygromycinR we replaced Blasticidin from the original vector (Addgene #52962) by HygromycinR using Gibson Assembly. For generation of Lenti-EF1α-Ovalbumin-P2A-mCherry-P2A-BlasticidinR, we replaced Cas9 (Addgene #52962) by Ovalbumin-P2A-mCherry derived from the original vector (Addgene #113030) with Gibson Assembly. The LC3-GFP-LC3ΔG-RFP construct was obtained from Addgene (#84572) and cloned into Lenti-EF1α-Cas9-P2A-BlasticidinR by replacing Cas9-P2A-BlasticidinR. The following vectors LentiGuide-Puro (#52963), LentiCRISPRv2-puro (#98290) and pLenti-PGK-Hygro-KRAS(G12V) (#35635) were obtained from Addgene.

### Lentivirus production

For lentivirus production of CRISPR libraries, transgenes or single guide HEK293T cells were transfected with, HEK293T cells (ATCC) were seeded in T175 cell culture flasks in DMEM (Gibco) supplemented with 10% FBS and 1% Penicillin/Streptomycin and grown up to 70% confluency. HEK293T cells were transfected with the following mixture: 10.4 *μ*g psPAX2-Plasmid, 3.5 *μ*g pMD2.G, 13.8 *μ*g lentiviral vector of interest (see section “Plasmids”) in a volume of 1000 *μ*l Opti-MEM (Gibco) in tube 1. In a second tube, 138 *μ*l 1mg/mL PEI (Polysciences) was mixed with 862 *μ*l Opti-MEM. Both tubes were incubated at room temperature for 5 min, mixed, and incubated again 20 min at room temperature and added to the cells in the evening. The next morning, medium was refreshed. After 48h and 72h, the supernatant was harvested, filtered with 0.45 *μ*m syringe filters (Sarstedt) and concentrated by centrifugation at 24,000 g for 2h. Plasmids psPAX2 (Addgene #12260) and pMD2.G (Addgene #12559) were gifts from Didier Trono.

### Genome wide CRISPR screen

The genome-wide Brie CRISPR-KO library (4 sgRNAs per gene; ~ 80.000 sgRNAs) was purchased from Addgene (#73632) and amplified according to the supplier’s protocol. KPC-Cas9-OVA were infected with the lentiviral Brie CRISPR-KO library at a multiplicity of infection (MOI) of 0.3 while keeping a 500x coverage of the library. One day post transduction, cells were selected with 2 *μ*g/ml Puromycin for 5 days in order to select for successfully transduced cells and to allow gene CRISPR-mediated gene knockout. For OT-I T-cell co-culture, 4×10^7^ KPC cells were plated at a final confluency around 60% and incubated for 3 days together with activated T-cells at an E:T ratio of 1:1. After T-cell killing, surviving KPC cells were left to recover for another 3 days. Untreated KPC-Cas9-OVA-Brie cells were cultured alongside and harvested for DNA isolation together with OT-I treated cells. For each replicate and sample, DNA was isolated from 4×10^7^ KPC cells using the Blood & Cell Culture DNA Maxi Kit (Qiagen). NGS libraries were prepared using the following primers:

### Staggered P5 forward primer

5’AATGATACGGCGACCACCGAGATCTACACTCTTTCCCTACACGACGCTCTTCCGATCT[N_1-8_]T TGTGGAAAGGACGAAACACCG

### Barcoded P7 reverse primer

5’CAAGCAGAAGACGGCATACGAGATNNNNNNNNGTGACTGGAGTTCAGACGTGTGCTCTT CCGATCTTCTACTATTCTTTCCCCTGCACTGT For each sample, total DNA obtained from 4×10^7^ cells was used as input with 10 *μ*g DNA per 100 *μ*l PCR reaction. DNA was amplified using Herculase II Fusion DNA Polymerase (Agilent) according to the manufacturer’s conditions with 2 *μ*l Herculase II and 2.5 mM MgCl_2_. Annealing was performed at 55°C and a total of 24 cycles. PCR reactions were cleaned up using 0.8x AMPure beads (Beckman Coulter). NGS libraries were run on the Illumina NovaSeq 6000 System generating 100 bp single-end reads.

### Data analysis

Demultiplexed reads were trimmed to exact 20 bp (sgRNA) using cutadapt. Subsequently, read counts were assed using MAGeCK (v0.5.6) as well as sgRNA enrichment/depletion with “mageck test -k *readcouts.txt* -t OT-I -c CTRL --norm-method control --control-sgrna *nontargeting*.txt”. For MAGeCK analysis three biological screening replicates were pooled. For pathway analysis of gene enrichment/depletion the cut-off was set at FDR < 0.1 and GO term analysis for candidates was performed using the Molecular Signature Database (MSigDB). The screening data set can be found in Tables S1 and S2 and via the GEO accession number GSE180834.

### Isolation and activation of CD8 T-cells

CD8^+^ OT-I T-cells were isolated from spleen, axillary and inguinal lymph nodes from OT-I mice. CD8^+^ cells were enriched using magnetic beads for MACS (130-104-075, Milteny Biotec). T-cells were cultured in IMDM supplemented with 10% fetal bovine serum (FBS), 1% penicillin/streptomycin (Gibco), 50 *μ*M β-Mercaptoethanol (Gibco) and 100 ng/ml Il-2 (Peprotech). Cells were kept at 37°C in 5% CO_2_. After T-cell isolation, cells were activated for 24h using 2 *μ*g/ml of anti-Cd28 and anti-Cd3ε antibodies (102116 and 100340, BioLegend).

### In vitro cytotoxicity assays

One day prior to OT-I co-culture, a total of 5×10^4^ Ovalbumin-expressing KPC cells were plated into 24-well plates. For competition assays, KPC-Cas9-OVA-mCherry (with CRISPR-KO of indicated gene) and KPC-Cas9-OVA-EGFP-sgRNA@ctrl were plated in a 1:1 ratio. Activated OT-I T-cells were added in the presence of 100 ng/ml Il-2 for one to two days, depending on downstream analysis. EGFP:mCherry ratio was assessed using flow cytometry. In brief, cells were trypsinized, spun down and washed in FACS buffer (2% FBS, 2mM EDTA in PBS). For MHC class I assessment, cells were detached using 5 mM EDTA. The following antibodies were used: CD8a-APC (1:600), CD8a-FITC (1:600), H-2Kb-APC (1:100), SIINFEKL-H-2Kb-APC (1:100; all Biolegend). SYTOX Blue was used as viability dye. For intracellular Granzyme-B staining (GzmB-FITC; GB11; BioLegend) cells were stained with eFluor780 (fixable viability dye) and stained with 4% formalin before permeabilization. GzmB-FITC staining was carried out in permeabilzation buffer for 30 min at room temperature. The following sgRNAs were cloned into the LentiGuide-puro vector and stably integrated as a pool per gene into target cells: *Ctrl-1*: 5’ GCGAGGTATTCGGCTCCGCG, *Rnf31-1*: 5’ CTACCTCAACACCCTATCCA, *Rnf31-2*: 5’ CCACCGTGCTGCGAAAGACG, *Rnf31-3*: 5’ TTCACTGAGCGCCAATACCG, *Vps4b-1*: 5’ TAAAGCCAAGCAAAGTATCA, *Vps4b-2*: 5’CGATAGAGCAGAAAAACTAA, *Vps4b-3*: 5’ TCAGGCCCAGTTGATGAGAA, *Stat1*: 5’ GGATAGACGCCCAGCCACTG, *Atg5*: 5’ AAGAGTCAGCTATTTGACGT.

### Autophagy flux assay

LC3-GFP-LC3ΔG-RFP construct was stably integrated into KPC-Cas9-OVA cells and GFP^+^/RFP^+^ single cell clones were sorted and expanded. Cells were starved for 8 hours in 2% FBS/PBS at 37°C in ambient CO_2_. Subsequently, cells were collected for flow cytometry analysis in order to assess GFP and RFP expression. The autophagic flux was assessed by calculating the GFP/RFP ratio and comparison to the non-starved control condition.

### Human pancreatic cancer organoids

Normal human pancreatic organoids (hPan) and pancreatic cancer organoids (hPDA) were generated as described elsewhere ^39^. Expansion medium (EM) contained Advanced DMEM/F12 supplemented with 10 mM HEPES, 1x Glutamax, 1% Penicillin/Streptomycin, 1x B27 without vitamin A (all Gibco), 1.25 mM N-acetylcysteine (Sigma), 25% WNT3A-conditioned medium (CM), 10% RSPO1-CM, 10% NOGGIN-CM (all CM produced in-house), 10 mM Nicotinamide (Sigma), 50 ng/mL human EGF (Peprotech), 100 ng/ml FGF10 (Peprotech), 10 nM Gastrin (Tocris Bioscience), 0.5 *μ*M TGF-b type I receptor inhibitor A83-01 (Tocris Bioscience) and 1*μ*M PGE2 (Tocris Bioscience). Organoids were split every 7-10 days using TrypLE Express (Gibco) and fire-polished Pasteur pipettes in a 1:3 – 1:4 ratio. After passaging, organoids were plated in 20 *μ*l drops of Matrigel (Corning) and overlaid with EM supplemented having 10 *μ*M RhoKinase inhibitor (Y-27632; Abmole). Organoids from healthy donors were lentivirally transduced to express oncogenic *KRAS*^G12V^ and to knockout *TP53* (hPan-KP). Both, hPan-KP and hPDA organoids were transduced with LentiCRISPRv2-puro to knockout *RNF31*. For lentiviral transduction, 3 – 4 full drops of organoids per condition were processed into single cells, mixed with 500 *μ*l EM + 10 *μ*M RhoKinase inhibitor + 50 *μ*l concentrated virus and spun for one hour at 32°C at 300 g in a 24-well plate. After 3 – 4 hours incubation at 37°C, cells were collected and plated in Matrigel. Organoid were selected with 1.5 *μ*g/ml puromycin (*RNF31*-KO), 10 *μ*M Nutlin-3a (*TP53*-KO; Sigma) and 300 *μ*g/ml hygromycin (Kras^G12V^). The following sgRNAs were used - *TP53*: 5’ GAAGGGACAGAAGATGACAG, *RNF31-1*: 5’ CCACCGTGCTGCGAAAGACA, *RNF31-2*: 5’ CCCAACCCCTTACAGCCTCG, *RNF31-3*: 5’ GGATCATGCTCACTAGCTGG.

### Sublibrary generation

For sublibrary generation, the top candidates (enriched and depleted; FDR < 10%) were selected (Table S3). If there were many genes within one pathway, only a couple of genes was selected to avoid redundancy. For each gene of the gene list (63 candidates and Ovalbumin) 10 sgRNAs (7 sgRNAs for Ovalbumin) were designed with the GPP sgRNA designer (Broad Institute). A total of 600 non-targeting controls was likewise included. Oligonucleotides having BsmBI (EspI) restriction sites, a single guide RNA sequence as well as primer binding sites for oligo pool amplification were synthesized (Twist Bioscience). Oligo sequences can be found in Table S3. PCR amplification of the oligo pool prior to cloning was done according to the manufacturer’s protocol. For cloning the oligo pool into the appropriate lentiviral backbone the following reaction was set up: 5 *μ*l 10x Cutsmart buffer (NEB), 1 mM DTT (final), 1 mM ATP (final), 1.5 *μ*l T4 DNA Ligase (8000U, NEB), 1.5 *μ*l EspI (NEB), 100 ng oligo pool PCR product and 500 ng vector (LentiGUIDE-puro, EspI-digested and isopropanol purified) and water up to 50 *μ*l. The reaction was incubated for 100 cycles at 5 min 37°C followed by 5 min 20°C. After isopropanol clean-up, the ligation was transformed into NEB Stable Competent *E. coli* (C3040I) and streak out onto LB agar plates. Library integrity was confirmed using Illumina sequencing.

### In vivo sublibrary screen

KPC-Cas9-OVA cells were transduced with the lentiviral sublibrary at a MOI of 0.3 and selected for four days with 2 *μ*g/ml Puromycin. Subsequently, 150.000 KPC-Cas9-OVA-Sublibrary cells were orthotopically transplanted into Rag1^−/−^ mice. On day 16 post transplantation 1 × 10^6^ preactivated OT-I CD8^+^ T-cells were adoptively transferred (intravenously) into tumor bearing mice. Mice were sacrificed on day 21 post transplantation and tumors were harvested. Tumor DNA was isolated using the Qiagen Blood and Tissue Kit and sgRNA cassette was amplified similarly to the *in vitro* screen and analyzed by Illumina sequencing. Single guide RNA representation was assed using MAGeCK (v0.5.6) by comparison to the plasmid sublibrary. The screening data set can be found in Table S3 and via the GEO accession number GSE180834.

### Transplantations

Mice were anesthetized using isoflurane at a constant flow rate. Abdomen was shaved and sterilized before small incision in the upper left quadrant was made. The pancreas was carefully put onto a cotton-swab and 1.5 ×10^5^ KPC cells were injected in 50 *μ*l of PBS:Matrigel (1:1) using a 29G needle. Successful injection was confirmed when a liquid bled formed and no leakage could be observed. Peritoneum and skin were subsequently sutured with Vicryl violet sutures (N385H, Ethicon) and secured with wound clips (FST). Approximately three weeks post transplantations animals were sacrificed and tumors were isolated, weighed and processed for subsequent analysis. For subcutaneous transplantations, C57BL/6 mice were injected with 1×10^6^ KPC or KPC-Cas9 cells per flank mix in 1:1 PBS:Matrigel. Tumors were measured with calipers and the volume was estimated via the equation (L×W^2^)/2.

### Flow cytometry for tumor microenvironment analysis

For flow cytometry analysis orthotopic tumors were collected and minced into small pieces before digestion in Collagense IV (6000 U/ml) and DNase I (200 U/ml) for one hour at 37°C. Cell suspension was filtered through a 40 *μ*m cell strainer. For cytokine stainings, cells were restimulated with PMA (100 nM), ionomycin (1 *μ*g/ml) and monensin (2 *μ*g/ml) for 3-4 hours at 37°C in complete IMDM medium (Gibco). Viability staining was performed using the fixable viability dye eFluor780. Staining with fluorescent antibodies was carried out for 15 min at 4°C in the dark. After washing in FACS buffer (2 mM EDTA, 2 % FBS), cell suspension was acquired using BD Fortessa and FlowJo software (Treestar).

### Antibodies used for flow cytometry

PD-1 FITC (J43; eBioscience), NK1.1 PE (PK136; eBioscience), CD3e PE-Dazzle594 (145-2C11; BioLegend), CD3e PE (145-2C11; eBioscience), FoxP3 PerCP-Cy5.5 (FJK-16s; eBioscience), CD8a PE-Cy7 (53-6.7; eBioscience), CD8a APC (53-6.7; eBioscience), CD8a PerCP-Cy5.5 (53-6.7; eBioscience), TCRb AF700 (H57-597; BioLegend), CD45 BV785 (30-F11; BioLegend), CD4 BV711 (GK1.5; BioLegend), CD19 BV650 (6D5; BioLegend), CD19 PE (1D3; eBioscience), CD11b BV605 (M1/70; BioLegend), CD11b BV510 (M1/70; BioLegend), CD11c BV605 (N418; BioLegend), Siglec-F PE (E50-2440; BD Biosciences), F4/80 APC (BM8; BioLegend), Ly-6G AF 700 (1A8; BioLegend), MHC II BV650 (M5/114.15.2; BioLegend), CD64 BV421 (X54-5/7.1; BioLegend), TNF FITC (MP6-XT22; BioLegend), IFNg PE-Cy7 (XMG1.2; BD Biosciences), GzmB FITC (GB11; BioLegend). Gating strategy is depicted in Supplementary Figure 6d.

### RNA-Seq

After 6h of co-culture, OT-I T-cells and KPC-Cas9-OVA cancer cells were sorted using the BD Aria cell sorter. KPC cells expressed mCherry, T-cells were stained with CD8a-APC (clone 53-6.7; Biolegend 100711) to separate both populations. RNA was isolated with the Qiagen RNeasy Mini Kit and sent to the Functional Genomic Center Zurich (FGCZ) for standard library preparation and Illumina sequencing. Reads were trimmed using cutadapt and mapped to the mouse genome GRCm38 with HISAT2 followed by sorting using samtools. The raw count matrix was generated in RStudio using Rsubread. Differential gene (DE) expression analysis was performed with EdgeR and differentially expressed genes (LFC ± 1; FDR < 0.1) were used as input for GO term analysis using the Molecular Signature Database (MSigDB). RNA-Seq data can be accessed via GEO.

### TNF treatment of KPC cells

KPC cells were treated 24h with the indicated TNF concentration and stained subsequently for immunofluorescence (see below) or with crystal violet to assess cell viability (here in the presence of 1 *μ*g/ml Actinomycin D). Crystal violet dye was reconstituted in 10% acetic acid and absorbance was measured at 595 nm.

### Immunofluorescence of KPC cells

Cells were grown on glass cover slips, treated with TNF (100 ng/ml) and fixed for 10 minutes at room temperature in 4% PFA. Cells were permeabilized and blocked in 0.5% Triton-X, 5% normal donkey serum in PBS. Cleaved caspase 3 antibody (1:400, Cell signaling Technology, 9664) was diluted in blocking solution and incubated overnight at 4°C. Coverslips were washed in PBS and incubated 2h at room temperature with secondary antibody (Donkey anti-rabbit-568, ThermoFisher Scientific, 1:400) and DAPI. Coverslips were mounted with Prolong Gold (ThermoFisher Scientific) and imaged with a Lunaphore.

### Whole mount staining of human pancreatic cancer organoids

Wildtype or *RNF31*^KO^ hPan/hPDA organoids were treated for 4h with 100 ng/ml human TNF (Peprotech) in 8-well *μ*-slides (Ibidi). After fixation in 4% PFA, organoids were blocked and permeabilized in blocking solution (10 % normal donkey serum; 0.5% Triton-X in PBS). All antibody incubations were performed overnight at 4°C on a rocking platform. Primary antibodies: E-Cadherin (1:500, R&D Systems, AF748), cleaved Caspase 3 (1:400, Cell Signaling Technology, 9664). Donkey-anti-goat 488 and donkey-anti-rabbit 568 were used as secondary antibodies and counterstained with DAPI. Organoids were mounted with ProLong Gold. Confocal Images were taken with a Zeiss LSM 880 Airyscan.

### Western Blot

Whole cell lysates were prepared in RIPA buffer (50mMTris-HCl pH 8.0, 150mMNaCl, 0.1% SDS, 0.5% Na-Deoxycholate, 1%IGEPAL CA-630) supplemented with PhosSTOP phosphatase inhibitors and cOmplete protease inhibitor cocktail (both Roche). BCA protein assay (ThermoScientific) was used for protein quantification. Samples were loaded on 4%– 15% precast polyacrylamide gels (Bio-Rad) and transferred to PVDF (Bio-Rad) membranes in Towbin buffer. Membranes were blocked in 5% Bovine Serum Albumin (Applichem) and incubated overnight in primary antibodies phospho-p65 (1:1000; CST#3033), Caspase 8 (1:1000; CST#4790), cleaved Caspase 8 (1:1000, CST#8592) and Gapdh (1:3000; CST#14C10). IRDye800CW and 680RD donkey anti-rabbit secondary antibodies were used for detection (LI-COR). Protein bands were visualized with the ODYSSEY CLx imaging system (LI-COR).

### Statistics

To compare data from experiments we either applied student’s unpaired, two-tailed T-test or One-Way ANOVA analysis, as indicated in the respective figure legend. A minimum of three independent biological replicates was performed per experiment. P values less than 0.05 were considered significant and significance levels were set as follows: *p < 0.05, **p < 0.01, ***p < 0.001. Statistical comparisons were performed using RStudio and GraphPad Prims 9.

## Data and materials availability

RNA-Seq data and CRISPR screening data have been made accessible via GEO: GSE180834. KPC cell line was received from Ximbio (Cat# 153474), including a materials transfer agreement.

## Acknowledgement

We thank the Flow Cytometry Facilities of the University of Zurich and the ETH Zurich. Also, we thank the Functional Genomics Center Zurich and the ETH Phenomics Center for their support and infrastructure.

## Author Information

N.F. and G. S. conceptualized the study, performed experiments, analyzed the data and wrote the manuscript. L.T. designed, supervised and analyzed flow cytometry experiments and gave valuable input throughout the course of the project. S.J. helped analyzing RNA-Seq data. D.E., K.F.M., T.R. and F.A. performed experiments. N.F. and G.S. wrote the manuscript, L.T. and M.K. reviewed and edited the manuscript. G.S. supervised the study. G.S. and M.K. acquired funding. All authors approved the final version of the manuscript.

## Supplementary Information

Supplementary Figures 1 – 6

Supplementary Tables 1 – 3

Supplementary References

