## Supplementary Figures and Tables for "Loss of Rnf31 and Vps4b sensitizes pancreatic cancer to T cell-mediated killing"

**This PDF file includes:**

Supplementary Figures 1 – 6

Supplementary Tables 1 – 3

Supplementary References

### Supplementary Figures

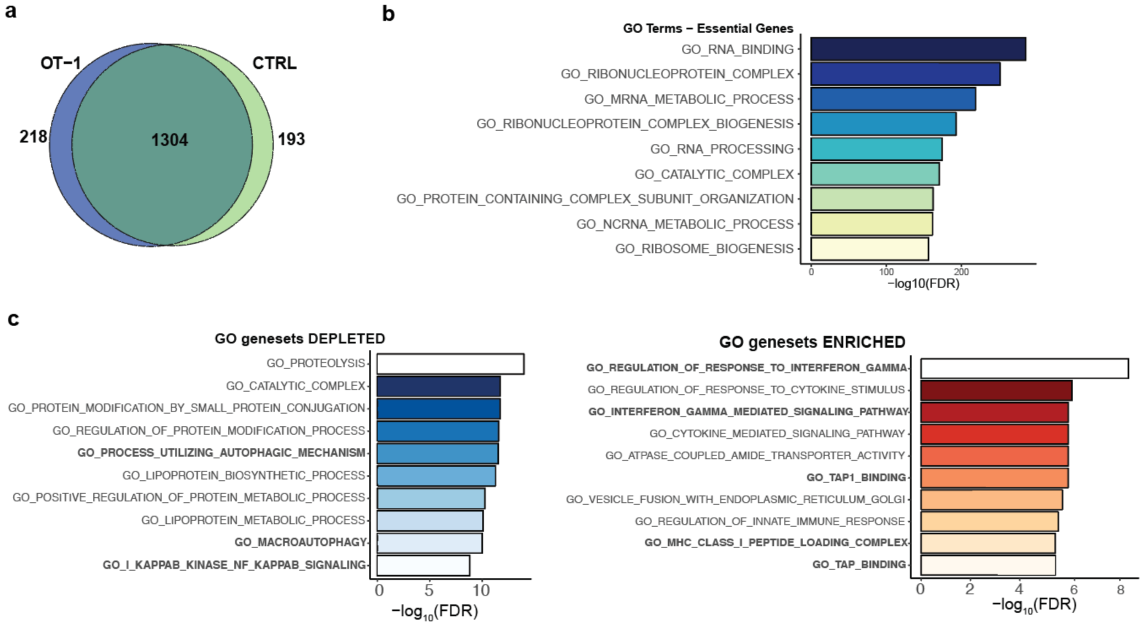


**Supplementary Figure 1. Related to Figure 1: Genome-wide CRISPR screen unravels immune evasion mechanisms in PDA. (a)** Venn diagram of essential genes in KPC-OVA cells untreated (CTRL) or after 3d OT-1 T cell treatment (OT-I). Guide RNA abundance was compared to plasmid Brie library using MAGeCK to find essential genes. **(b)** Pathway analysis of significantly depleted genes in CTRL and OT-I conditions (1304 genes, see S1A) using the Molecular Signature Database (MSigDB). **(c)** Pathway analysis of significantly enriched/depleted genes in OT-I treated vs. untreated KPC cells (FDR < 0.1) using the Molecular Signature Database (MSigDB). The -log10 FDR of the top ten pathways are represented

**a**

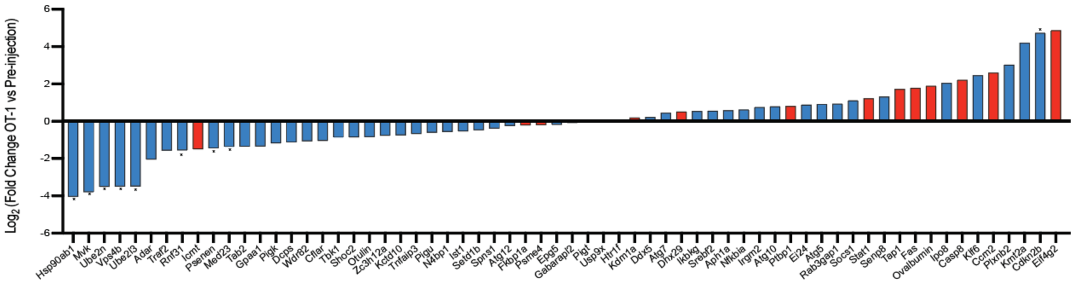


**Supplementary Figure 2: Related to Figure 2: In vivo CRISPR screening validates hits identified in the in vitro CRISPR screen. (a)** $Log_2$Fold change of all sublibrary genes (OT-I treated mice vs. pre-injection pool of cells using MAGeCK). Red and blue indicates predicted to be enriched or depleted based on genome-wide in vitro screen, respectively. Asterisks indicate FDR < 0.2.

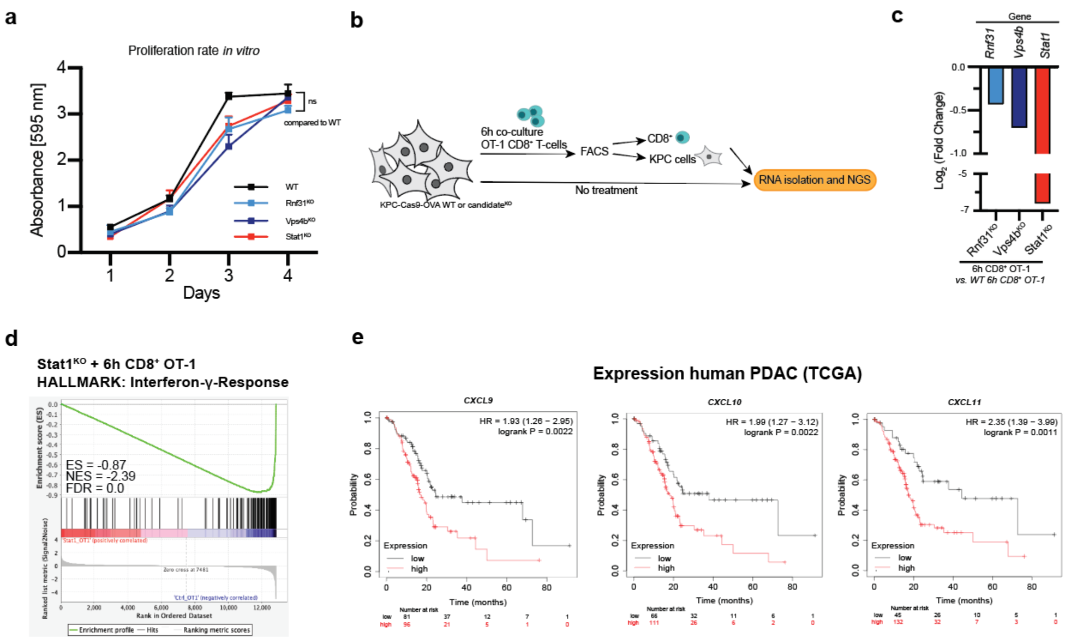


**Supplementary Figure 3. Related to Figure 3: *Rnf31*$^{KO}$ and *Vps4b*$^{KO}$ alter transcriptional response upon T cell exposure.** **(a)** Proliferation rate of different cell lines. Crystal violet staining was carried out and normalized to day 1. Significance was determined with an unpaired two-tailed t test. *p < 0.05, **p < 0.01, ***p < 0.001; ns, non-significant, p > 0.05. Error bars represent ± SD. **(b)** Schematic of workflow for RNA Sequencing sample generation. **(c)** $Log_2$ fold change of candidate genes compared to KPC control cells. **(d)** Gene Set Enrichment Analysis (GSEA) of *Stat1*$^{KO}$ cells to control KPC cells after T cell exposure. **(e)** Kaplan-Meier plot of human pancreatic cancer cases (n = 177) from the cancer genome atlas (TCGA) analyzed according to the platform “Kaplan-Meier Plotter” [2].

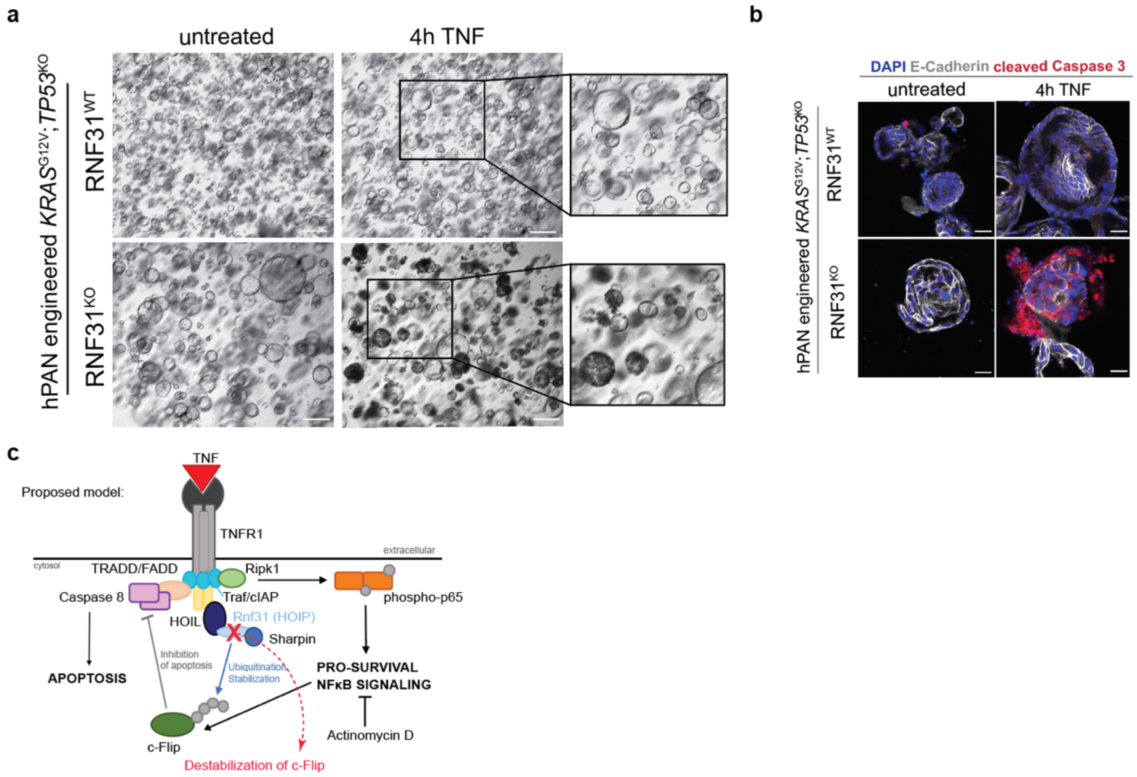


**Supplementary Figure 4. Related to Figure 4: *Rnf31*$^{KO}$ sensitizes PDA to TNF-induced apoptosis in human 3D organoids. (a)** Brightfield images of human engineered pancreatic organoids in the presence of 100 ng/ml TNF for 4h. Boxes highlight viable and dying organoids. Scale bar represents 200 $\mu$m. **(b)** Whole mount staining of engineered human pancreatic organoids after 4h TNF (100 ng/ml) treatment with cleaved caspase 3 (red), E-cadherin (white) and DAPI (blue). Scale bar represents 20 $\mu$m. **(c)** Schematic of TNF-induced signaling cascade with emphasis on LUBAC. Adapted from Tang *et al*. 2018 [3].

a

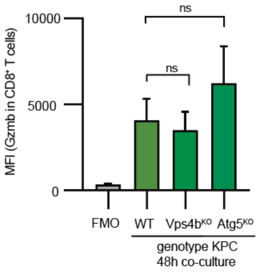


**Supplementary Figure 5. Related to Figure 5:** ***Vps4b^KO*** **abrogates autophagy and increases granzyme B levels. (a)** Mean fluorescence intensity (MFI) of granzyme B contents in CD8$^+$ OT-I T cells. FMO (Fluorescence Minus One) control for Gzmb-FITC antibody. Significance was determined with one-way ANOVA. *$p < 0.05$, **$p < 0.01$, ***$p < 0.001$; ns, non-significant, $p > 0.05$. Values represent mean ± SEM, data are derived from at least three independent experiments.

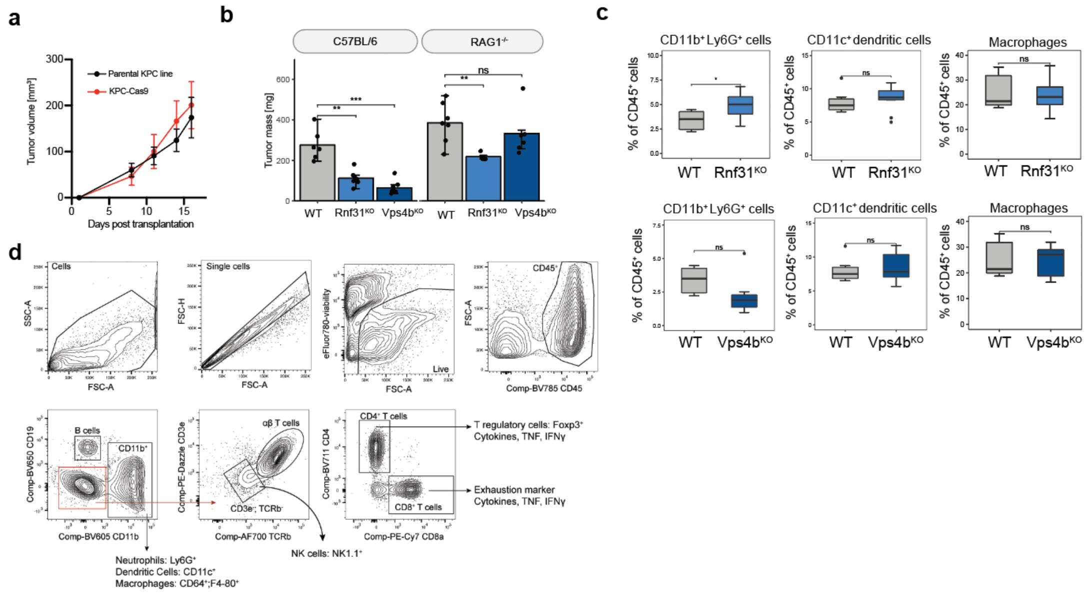


**Supplementary Figure 6. Related to Figure 6:** ***Rnf31*$^{KO}$ and *Vps4b*$^{KO}$ alter tumor growth *in vivo*.** **(a)** Tumor growth of subcutaneous KPC (black) and KPC-Cas9 (red) tumors into C57BL/6 mice. **(b)** Tumor mass of candidate KPC line in immune competent C57BL/6 mice and RAG1$^{-/-}$ mice. **(c)** Flow cytometry analysis of immune cell population within tumors. **(d)** Gating strategy for TME characterization. Significance in **(a)** and **(c)** was determined with an unpaired two-tailed t test. *$p < 0.05$, **$p < 0.01$, ***$p < 0.001$; ns, non-significant, $p > 0.05$. Values represent mean ± SD.

**Supplementary Table 1:** Gene summary (MAGeCK RRA) of top candidate genes in genome-wide OT-I screen (FDR < 0.1). OT-I treated vs untreated KPC cells.

| | | Positive Selection | | Negative Selection | |
|---|---|---|---|---|---|
| **Gene** | **LogFC** | **p value** | **FDR** | **p value** | **FDR** |
| Ifngr2 | 2.2254 | 2.40E-07 | 0.00055 | 1 | 1 |
| Ifngr1 | 2.3361 | 2.40E-07 | 0.00055 | 1 | 1 |
| Stat1 | 2.2376 | 2.40E-07 | 0.00055 | 1 | 1 |
| Jak1 | 2.204 | 2.40E-07 | 0.00055 | 1 | 1 |
| Jak2 | 1.6873 | 2.40E-07 | 0.00055 | 1 | 1 |
| Ccm2 | 0.93447 | 2.40E-07 | 0.00055 | 1 | 1 |
| Kdm1a | 0.95777 | 2.40E-07 | 0.00055 | 1 | 1 |
| Casp8 | 1.0021 | 2.40E-07 | 0.00055 | 1 | 1 |
| Fkbp1a | 1.0791 | 2.40E-07 | 0.00055 | 1 | 1 |
| Ptbp1 | 0.83122 | 4.07E-06 | 0.007651 | 0.99986 | 1 |
| Tap1 | 0.83631 | 4.07E-06 | 0.007651 | 0.99999 | 1 |
| Icmt | 0.69433 | 5.51E-06 | 0.009488 | 1 | 1 |
| Eif4g2 | 0.62934 | 2.08E-05 | 0.032885 | 0.99986 | 1 |
| Tapbp | 0.71959 | 2.23E-05 | 0.032885 | 0.99999 | 1 |
| Tap2 | 0.65265 | 3.23E-05 | 0.044554 | 0.99999 | 1 |
| Fas | 0.69197 | 3.66E-05 | 0.047339 | 0.99913 | 1 |
| Psme4 | 0.7215 | 6.78E-05 | 0.082411 | 0.99993 | 1 |
| Rnf31 | -1.3877 | 1 | 1 | 2.40E-07 | 0.000381 |
| Vps4b | -1.2716 | 1 | 1 | 2.40E-07 | 0.000381 |
| Gabarapl2 | -0.86886 | 1 | 1 | 2.40E-07 | 0.000381 |
| Klf6 | -0.62437 | 1 | 1 | 2.40E-07 | 0.000381 |
| Tnfaip3 | -0.95731 | 1 | 1 | 2.40E-07 | 0.000381 |
| Gpaa1 | -1.0891 | 1 | 1 | 2.40E-07 | 0.000381 |
| Ube2n | -1.1611 | 1 | 1 | 2.40E-07 | 0.000381 |
| Cflar | -0.91552 | 1 | 1 | 2.40E-07 | 0.000381 |
| Atg5 | -1.115 | 0.99997 | 1 | 2.40E-07 | 0.000381 |
| Aph1a | -0.85653 | 0.99634 | 1 | 2.40E-07 | 0.000381 |
| Adar | -1.1541 | 0.99829 | 1 | 2.40E-07 | 0.000381 |
| Ei24 | -1.1046 | 0.99973 | 1 | 2.40E-07 | 0.000381 |
| Srebf2 | -0.93756 | 0.99691 | 1 | 2.40E-07 | 0.000381 |
| Irgm2 | -0.92123 | 1 | 1 | 7.19E-07 | 0.00099 |
| Shoc2 | -0.83244 | 1 | 1 | 7.19E-07 | 0.00099 |
| Tab2 | -1.0876 | 0.99973 | 1 | 1.20E-06 | 0.001456 |
| Traf2 | -0.8403 | 0.99996 | 1 | 1.20E-06 | 0.001456 |
| Nfkbia | -0.60428 | 1 | 1 | 2.87E-06 | 0.0033 |
| Usp9x | -0.71683 | 1 | 1 | 4.07E-06 | 0.004429 |
| Pigk | -0.58995 | 1 | 1 | 4.55E-06 | 0.004479 |
| Ube2l3 | -0.95439 | 1 | 1 | 4.55E-06 | 0.004479 |
| Kctd10 | -0.65842 | 1 | 1 | 5.51E-06 | 0.005176 |
| Tbk1 | -0.82417 | 0.90318 | 1 | 5.99E-06 | 0.005381 |
| Cdkn2b | -0.43096 | 1 | 1 | 6.95E-06 | 0.005982 |
| Med23 | -0.79003 | 1 | 1 | 7.43E-06 | 0.006139 |
| Epg5 | -0.69529 | 0.79229 | 1 | 8.38E-06 | 0.006664 |
| Atg12 | -0.9694 | 0.95429 | 1 | 9.34E-06 | 0.007151 |
| Ddx5 | -0.87276 | 1 | 1 | 9.82E-06 | 0.007249 |
| Socs1 | -0.96168 | 0.83603 | 1 | 1.27E-05 | 0.009047 |
| Atg10 | -0.69371 | 1 | 1 | 1.65E-05 | 0.011386 |
| Ist1 | -0.79476 | 0.99131 | 1 | 1.94E-05 | 0.012935 |
| N4bp1 | -0.5666 | 0.99999 | 1 | 3.09E-05 | 0.019652 |
| Pigt | -0.76161 | 0.99992 | 1 | 3.14E-05 | 0.019652 |
| Setd1b | -0.61491 | 0.99999 | 1 | 3.23E-05 | 0.019656 |
| Spns1 | -0.4862 | 0.99998 | 1 | 5.01E-05 | 0.029562 |

| | | | | | |
|---|---|---|---|---|---|
| Kmt2a | -0.53789 | 0.99998 | 1 | 5.58E-05 | 0.032041 |
| Atg7 | -0.67195 | 0.22399 | 0.998633 | 6.30E-05 | 0.035189 |
| Zc3h12a | -0.53973 | 0.99974 | 1 | 8.74E-05 | 0.047551 |
| Hsp90ab1 | -0.94902 | 0.90062 | 1 | 9.41E-05 | 0.049886 |
| Plxnb2 | 0.33279 | 0.017553 | 0.811154 | 9.99E-05 | 0.051609 |
| Ipo8 | -0.47609 | 0.92711 | 1 | 0.00012048 | 0.060734 |
| Ikbkg | -0.62624 | 0.85092 | 1 | 0.00012719 | 0.062514 |
| Pigu | -0.66735 | 0.99982 | 1 | 0.00013006 | 0.062514 |
| Psenen | -0.6673 | 0.99203 | 1 | 0.00015833 | 0.07437 |
| Rab3gap1 | -0.44779 | 0.8693 | 1 | 0.00016455 | 0.075578 |
| Mvk | -0.77305 | 0.99815 | 1 | 0.00018419 | 0.082759 |
| Senp8 | -0.54521 | 0.99176 | 1 | 0.00019473 | 0.085633 |
| Htr1f | -0.44009 | 0.99994 | 1 | 0.00021964 | 0.094575 |
| Otulin | -0.6721 | 0.99994 | 1 | 0.00023018 | 0.09709 |

**Supplementary Table 2:** List of genes in the targeted CRISPR sublibrary based on top candidates of the genome-wide *in vitro* screen (FDR < 0.1)

| Gene | Number of sgRNAs | Prediction GWS screen |
|---|---|---|
| Casp8 | 10 | Enriched |
| Ccm2 | 10 | Enriched |
| Eif4g2 | 10 | Enriched |
| Fas | 10 | Enriched |
| Fkbp1a | 10 | Enriched |
| Icmt | 10 | Enriched |
| Kdm1a | 10 | Enriched |
| Ovalbumin | 7 | Enriched |
| Psme4 | 10 | Enriched |
| Ptbp1 | 10 | Enriched |
| Stat1 | 10 | Enriched |
| Tap1 | 10 | Enriched |
| Adar | 10 | Depleted |
| Aph1a | 10 | Depleted |
| Atg10 | 10 | Depleted |
| Atg12 | 10 | Depleted |
| Atg5 | 10 | Depleted |
| Atg7 | 10 | Depleted |
| Cdkn2b | 10 | Depleted |
| Cflar | 10 | Depleted |
| Dcps | 10 | Depleted |
| Ddx5 | 10 | Depleted |
| Dhx29 | 10 | Depleted |
| Ei24 | 10 | Depleted |
| Epg5 | 10 | Depleted |
| Gabarapl2 | 10 | Depleted |
| Gpaa1 | 10 | Depleted |
| Hsp90ab1 | 10 | Depleted |
| Htr1f | 10 | Depleted |
| Ikbkg | 10 | Depleted |
| Ipo8 | 10 | Depleted |
| Irgm2 | 10 | Depleted |
| Ist1 | 10 | Depleted |
| Kctd10 | 10 | Depleted |
| Klf6 | 10 | Depleted |
| Kmt2a | 10 | Depleted |
| Med23 | 10 | Depleted |
| Mvk | 10 | Depleted |
| N4bp1 | 10 | Depleted |
| Nfkbia | 10 | Depleted |
| Otulin | 10 | Depleted |
| Pigk | 10 | Depleted |
| Pigt | 10 | Depleted |
| Pigu | 10 | Depleted |
| Plxnb2 | 10 | Depleted |
| Psenen | 10 | Depleted |
| Rab3gap1 | 10 | Depleted |
| Rnf31 | 10 | Depleted |
| Senp8 | 10 | Depleted |
| Setd1b | 10 | Depleted |
| Shoc2 | 10 | Depleted |
| Socs1 | 10 | Depleted |
| Spns1 | 10 | Depleted |

| | | |
|---|---|---|
| Srebf2 | 10 | Depleted |
| Tab2 | 10 | Depleted |
| Tbk1 | 10 | Depleted |
| Tnfaip3 | 10 | Depleted |
| Traf2 | 10 | Depleted |
| Ube2l3 | 10 | Depleted |
| Ube2n | 10 | Depleted |
| Usp9x | 10 | Depleted |
| Vps4b | 10 | Depleted |
| Wdr82* | 10 | Depleted |
| Zc3h12a | 10 | Depleted |
| Non-targeting | 600 | Depleted |
| **Total:** | **1237** | |

**Supplementary Table 3:** Gene summary (MAGeCK RRA) of sublibrary in vivo screen. OT-I treated mice vs. plasmid library.

| | | Positive Selection | | Negative Selection | |
|---|---|---|---|---|---|
| **Gene** | **LogFC** | **p value** | **FDR** | **p value** | **FDR** |
| Vps4b | -3.4933 | 0.57416 | 1 | 4.95E-06 | 0.001096 |
| Hsp90ab1 | -4.0424 | 0.96674 | 1 | 4.95E-06 | 0.001096 |
| Ube2l3 | -3.4897 | 0.89437 | 1 | 4.95E-06 | 0.001096 |
| Mvk | -3.7923 | 0.99984 | 1 | 1.49E-05 | 0.001974 |
| Ube2n | -3.5023 | 0.35716 | 1 | 1.49E-05 | 0.001974 |
| Rnf31 | -1.5502 | 0.9635 | 1 | 0.00019321 | 0.021382 |
| Psenen | -1.4586 | 0.98627 | 1 | 0.00023284 | 0.022086 |
| Dcps | -1.1319 | 0.99124 | 1 | 0.0021847 | 0.161915 |
| Med23 | -1.3727 | 0.44142 | 1 | 0.0021946 | 0.161915 |
| Adar | -2.042 | 0.9033 | 1 | 0.0044339 | 0.294408 |
| Tab2 | -1.3684 | 0.57417 | 1 | 0.023636 | 1 |
| Traf2 | -1.5665 | 0.46052 | 1 | 0.024498 | 1 |
| Icmt | -1.4927 | 0.24827 | 1 | 0.028411 | 1 |
| Pigk | -1.1718 | 0.73845 | 1 | 0.028441 | 1 |
| Cflar | -1.0461 | 0.55064 | 1 | 0.031998 | 1 |
| Gpaa1 | -1.3499 | 0.71803 | 1 | 0.039181 | 1 |
| Shoc2 | -0.85451 | 0.26075 | 1 | 0.05763 | 1 |
| Wdr82 | -1.0617 | 0.99689 | 1 | 0.067558 | 1 |
| Tbk1 | -0.87305 | 0.98235 | 1 | 0.072998 | 1 |
| Pigu | -0.60473 | 0.8428 | 1 | 0.07708 | 1 |
| Tnfaip3 | -0.67185 | 0.87792 | 1 | 0.11683 | 1 |
| Otulin | -0.84912 | 0.28649 | 1 | 0.14763 | 1 |
| Kctd10 | -0.74863 | 0.35715 | 1 | 0.1867 | 1 |
| Epg5 | -0.1858 | 0.28649 | 1 | 0.19712 | 1 |
| N4bp1 | -0.56002 | 0.50731 | 1 | 0.20553 | 1 |
| Setd1b | -0.47321 | 0.28649 | 1 | 0.21116 | 1 |
| Zc3h12a | -0.76704 | 0.86264 | 1 | 0.30423 | 1 |
| Ist1 | -0.53971 | 0.35716 | 1 | 0.30496 | 1 |
| Irgm2 | 0.75237 | 0.27347 | 1 | 0.33906 | 1 |
| Pigt | -0.038195 | 0.80842 | 1 | 0.34589 | 1 |
| Atg7 | 0.45524 | 0.2865 | 1 | 0.35474 | 1 |
| Nfkbia | 0.62486 | 0.55065 | 1 | 0.35567 | 1 |
| Psme4 | -0.20224 | 0.31057 | 1 | 0.3936 | 1 |
| Spns1 | -0.3847 | 0.64684 | 1 | 0.40478 | 1 |
| Usp9x | 0.0098218 | 0.68419 | 1 | 0.42424 | 1 |
| Ikbkg | 0.54739 | 0.64685 | 1 | 0.44576 | 1 |
| Atg12 | -0.24401 | 0.46052 | 1 | 0.54292 | 1 |
| Kdm1a | 0.19833 | 0.75799 | 1 | 0.55631 | 1 |
| Fkbp1a | -0.21733 | 0.44141 | 1 | 0.56106 | 1 |
| Srebf2 | 0.55871 | 0.12455 | 1 | 0.61809 | 1 |
| Gabarapl2 | -0.085073 | 0.80445 | 1 | 0.6301 | 1 |
| Rab3gap1 | 0.93038 | 0.46051 | 1 | 0.64833 | 1 |
| Htr1f | 0.072868 | 0.98078 | 1 | 0.64922 | 1 |
| Ddx5 | 0.22954 | 0.35716 | 1 | 0.80202 | 1 |
| Atg10 | 0.79672 | 0.73324 | 1 | 0.81679 | 1 |
| Dhx29 | 0.51731 | 0.87303 | 1 | 0.85282 | 1 |
| Ei24 | 0.88513 | 0.016294 | 1 | 0.88165 | 1 |
| Klf6 | 2.4688 | 0.26076 | 1 | 0.88862 | 1 |
| Socs1 | 1.1162 | 0.45111 | 1 | 0.9505 | 1 |
| Ovalbumin | 1.8922 | 0.35239 | 1 | 0.94762 | 1 |
| Tap1 | 1.7225 | 0.2865 | 1 | 0.9632 | 1 |
| Plxnb2 | 3.0225 | 0.18298 | 1 | 0.98228 | 1 |

| | | | | | |
|---|---|---|---|---|---|
| Aph1a | 0.58394 | 0.71802 | 1 | 0.98492 | 1 |
| Atg5 | 0.9242 | 0.56681 | 1 | 0.99067 | 1 |
| Casp8 | 2.216 | 0.27347 | 1 | 0.99158 | 1 |
| Eif4g2 | 4.8799 | 0.13904 | 1 | 0.9936 | 1 |
| Senp8 | 1.3182 | 0.35715 | 1 | 0.99397 | 1 |
| Ptbp1 | 0.81344 | 0.56681 | 1 | 0.99769 | 1 |
| Stat1 | 1.22 | 0.24826 | 1 | 0.99962 | 1 |
| Fas | 1.7888 | 0.26076 | 1 | 0.99972 | 1 |
| Ipo8 | 2.0565 | 0.18299 | 1 | 0.99973 | 1 |
| Ccm2 | 2.6031 | 0.2986 | 1 | 0.99997 | 1 |
| Cdkn2b | 4.7317 | 0.00013376 | 0.088816 | 1 | 1 |
| Kmt2a | 4.2092 | 0.047801 | 1 | 1 | 1 |
